# Complex coding and regulatory polymorphisms in a restriction factor determine the susceptibility of *Drosophila* to viral infection

**DOI:** 10.1101/117846

**Authors:** Chuan Cao, Rodrigo Cogni, Vincent Barbier, Francis M. Jiggins

## Abstract

It is common to find that major-effect genes are an important cause of variation in susceptibility to infection. Here we have characterised natural variation in a gene called *pastrel* that explains over half of the genetic variance in susceptibility to the virus DCV in populations of *Drosophila melanogaster*. We found extensive allelic heterogeneity, with a sample of seven alleles of *pastrel* from around the world conferring four phenotypically distinct levels of resistance. By modifying candidate SNPs in transgenic flies, we show that the largest effect is caused by an amino acid polymorphism that arose when an ancestral threonine was mutated to alanine, greatly increasing resistance to DCV. Overexpression of the ancestral susceptible allele provides strong protection against DCV, indicating that this mutation acted to improve an existing restriction factor. The *pastrel* locus also contains complex structural variation and *cis*-regulatory polymorphisms altering gene expression. We find that higher expression of *pastrel* is associated with increased survival after DCV infection. To understand why this variation is maintained in populations, we investigated genetic variation surrounding the amino acid variant that is causing flies to be resistant. We found no evidence of natural selection causing either recent changes in allele frequency or geographical variation in frequency, suggesting that this is an old polymorphism that has been maintained at a stable frequency. Overall, our data demonstrate how complex genetic variation at a single locus can control susceptibility to a virulent natural pathogen.

## Introduction

A central aim of infectious disease research is to understand why individuals within populations vary in their susceptibility to infection. This variation often has a substantial genetic component, and much effort has been devoted to identifying the genes involved (Cao, et al. 2016; Magwire, et al. 2011; Magwire, et al. 2012; Martins, et al. 2014). It is common to find that natural populations contain major-effect polymorphisms that affect susceptibility to infection, especially when natural pathogens or parasites are studied. In humans, for example, major-effect genes affect susceptibility to *Plasmodium falciparum* malaria, *Plasmodium vivax* malaria, HIV and Norwalk virus diarrhoea (Hill 2012). Studying these genes can not only advance our understanding of the mechanisms of resistance and functioning of immune systems, but it can also provide insights into evolutionary processes. For example, theoretical models of host-parasite coevolution make strong assumptions about the genetic basis of resistance (Routtu and Ebert 2015). More generally, pathogens are one of the most important selective agents in nature, so understanding the genetic basis of how host populations respond to this selection pressure is of great interest.

While much research has focussed on humans, crops and domestic animals, studying the natural pathogens of model organisms such as *Arabidopsis, Drosophila* and *C. elegans* provides a powerful way to understand the genetics of infectious disease resistance. The best-characterised natural pathogens of *Drosophila* are viruses, with most research focussing on the sigma virus (Rhabdoviridae; DMelSV (Longdon B 2012)) and Drosophila C virus (Dicistroviridae; DCV (Ferreira, et al. 2014; Hedges and Johnson 2008; Johnson and Christian 1998; Kemp, et al. 2013; Longdon, et al. 2013; Magwire, et al. 2012; Martins, et al. 2014; Zhu, et al. 2013)). DMelSV is a vertically transmitted virus that is relatively benign, causing a ~20% drop in fitness (Wilfert and Jiggins 2013; Yampolsky, et al. 1999). In contrast DCV is horizontally transmitted and multiplies in most tissues of adult *Drosophila melanogaster*, causing marked pathogenic effects and sometimes death (Chtarbanova, et al. 2014).

There is considerable genetic variation in susceptibility to both of these viruses within natural populations of *D. melanogaster* (Magwire, et al. 2012). Much of this variation is caused by major-effect polymorphisms that confer a high level of resistance. In the case of DMelSV, three polymorphic resistance genes have been identified—*p62 (ref(2)P)* (Bangham, et al. 2008; Contamine, et al. 1989), *CHKov1* (Magwire, et al. 2011) and *Ge-1* (Cao, et al. 2016). In a North American population *p62* and *CHKov1* together explain 37% of the genetic variance in susceptibility to DMelSV(Magwire, et al. 2012). Resistance to DCV is controlled by a very small number of genes, with a SNP in a gene called *pastrel* (*pst*) on chromosome III explaining 47% of the genetic variance in DCV susceptibility (Magwire, et al. 2012). In another mapping population of flies, we recently reported that this gene accounted for 78% of the genetic variance (Cogni, et al. 2016).

Despite its key role in virus resistance, *pst* remains poorly characterised. Its molecular function remains unknown, although it has been reported to participate in olfactory learning (Dubnau, et al. 2003), protein secretion (Bard, et al. 2006) and to be associated with lipid droplets (Beller, et al. 2006). We identified the gene using an association study on 185 lines from North America with complete genome sequences (Mackay, et al. 2012). In this study, 6 SNPs were found to be associated with resistance to DCV at P<10^-12^, including two adjacent SNPs in the 3’UTR (T2911C and A2912C), two non-synonymous SNPs (G484A and A2469G) and two SNPs in introns (C398A and A1870G). All of these are in linkage disequilibrium, and the non-synonymous SNP A2469G in the last coding exon stands out as the most significant polymorphism (Magwire, et al. 2012). However, the strong linkage disequilibrium between SNPs prevents us from identifying the causal SNP(s).

In this study, we have characterised genetic variation in *pst* and its effects on susceptibility to viral infection. In a sample of seven copies of the gene from natural populations, we find four functionally distinct alleles that confer varying levels of resistance. By combining association studies and transgenic techniques we identify an amino acid substitution that has led to a large increase in resistance. This appears to be a relatively old polymorphism that has been maintained at a relatively stable frequency in natural populations. The *pst* locus also contains complex structural variation and *cis* regulatory variation affecting gene expression. Higher levels of *pst* expression are associated with increased resistance. Therefore, this is a complex gene in which multiple genetic variants affecting both gene expression and the amino acid sequence alter susceptibility to viral infection.

## Materials and Methods

### Generating transgenic flies carrying alleles of pst modified by recombineering

To test which SNPs in *pst* are causing flies to be resistant, we used recombineering to modify a BAC (bacterial artificial chromosome) clone of the region of the *Drosophila* genome containing the gene (Warming, et al. 2005). This allowed us to make precise modifications of six candidate SNPs previously identified in *pst*, with five BACs carrying each SNP separately (SNP T2911C and SNP A2912C are adjacent and in complete linkage disequilibrium in nature, so were considered as single locus TA2911(2)CC).

*Drosophila* P[acman] Bacteria Artificial Chromosomes (BACs) were obtained from BACPAC Resources Centre (BPRC) (Venken, et al. 2009; Venken, et al. 2006). The *CHORI-322-21P14* clone, which covers a region of the fly genome that includes *pst* (genome positions: 3R:2114276-21164956), was chosen for its smaller size (20.064kb) and therefore higher transformation efficiency (Venken, et al. 2009). This BAC doesn’t contain any duplication or deletion of *pst*. BACs were extracted from original cells and transformed into sw102 cells. The sw102 strain was derived from the DY380 strain of *E.coli* with a deletion of the *galK* gene (Warming, et al. 2005). Competent sw102 cells were made following Warming’s protocol (Warming, et al. 2005).

In the BAC clone containing *pst*, we modified the candidate SNPs controlling resistance using recombineering and *GalK* positive-negative selection. This results in a ‘seamless’ modification, where the only difference between the two BAC clones is the SNP of interest. First, the selectable marker *GalK* was introduced to the site of the SNP, with positive selection for *GalK*. Second, *GalK* is replaced by the alternate allele of the SNP by selecting against *GalK*, resulting in a BAC clone that differs only in the nucleotide of interest. For the first step, *GalK* targeting cassettes were PCR-amplified from vector pgalK (Warming, et al. 2005) using five different pairs of primers, each of which has about 80 bp of sequence homologous to *pst* at each 5’ end. Phusion^®^ High Fidelity polymerase (NEB) was used in the following conditions: 95°C for 4 min, then 95°C for 15 s, 55°C for 30 s and 68°C for 1 min (1 min per 1 kb product), for 35 cycles, incubate at 68°C for 5 min. PCR products were gel-purified using Invitrogen PureLink^™^ Quick Gel Extraction Kit and always used freshly for transfection. Recombineering was carried out using protocol developed by Warming *et al*. (Warming, et al. 2005). The correct insertion of *GalK* was confirmed by PCR genotyping. The next step was to replace *GalK* with DNA fragment containing the SNP of interest. DNA was extracted from three DGRP lines (Mackay, et al. 2012) that had the desired sequence of *pst*. We then amplified a region of size ranging from 300bp to 1kb that contained the SNP of interest near the centre of the PCR product using the conditions described above. The PCR product was then purified and used to replace *GalK* using the protocol described above, this time selecting against *GalK* using 2-deoxy-galactose (DOG)(Warming, et al. 2005). The *pst* gene was then sequenced to check the process had been successful. Fly stocks used as template and primers used in PCR were listed in Table S1.

We next inserted the five modified BAC clones containing the different *pst* alleles into identical sites in the genome of a fly line. This was possible as the BACs contain an *attB* site, which allows them to be inserted into *attP* docking sites of flies (Bischof, et al. 2007). Plasmids of concentration between 0.1ug/ul and 0.3ug/ul and OD 260/280 ratio between 1.8-1.9 were injected into the embryos of an *attP* line: *y^−^w^−^M^(eGFP,vas-int,dmRFP)^ZH-2A;P{CaryP}attp40*. Male adults were crossed to a white-eye double balancer line *w^−^;If/Cyo;TM6B/MRKS*. The BAC contains a wild-type allele of *white*, allowing us to select for successful transformants by their red-eye phenotype. Male and female transformants were crossed to generate homozygotes with balanced third chromosomes. Homozygous transformants were then crossed to a balanced *pst* hypomorphic mutant *y^1^w^67c23^;If/Cyo;P^GSV1^GS3006/TM3,Sb^1^Ser^1^* (generated by crossing *pst* mutant *y^1^w^67c23^;P{GSV1}GS3006/TM3,Sb^1^Ser^1^* (DGRC #200404) to *w^−^;If/Cyo;TM6B/MRKS)* and generated *w;BAC^*^;P^GSV1^GS3006/TM3,Sb^1^Ser^1^*. This balanced hypomorphic mutant has a P-element inserted in the 5’UTR of the *pst* gene and has a lower *pst* mRNA expression level compared to many lab fly stocks we tested.

### Over-expressing pst in flies

Transgenic flies that overexpress two different *pst* alleles were generated using vector pCaSpeR-hs fused with *pst* sequence. Expression of *pst* was under the control of HSP70-promoter, and the protein is tagged by the FLAG epitope in N-terminus. The two *pst* alleles were amplified from cDNA from the fly lines DGRP-101 and DGRP-45 (Bloomington *Drosophila* Stock Center). These two DGRP lines encode identical *pst* amino acid sequences except for the Ala/Thr difference caused by SNP A2469G. Plasmids carrying different *pst* alleles were injected into a docker fly lines containing *attP* site on the second chromosome *y^−^ w^−^M^eGFP,vas-int,dmRFP^ZH-2A;P{CaryP}attp40*. The experiment was subsequently repeated a different fly line with a different *attP* site: *y^−^w^−^M^eGFP,vas-int,dmRFP^ZH-2A;M^attp^ZH-86Fb*. Male adults were crossed to white-eye balancer: *w^1118iso^/y^+^Y;Sco/SM6a;3^iso^* to select for successful transformants. Male and female transformants were crossed to generate homozygotes. The *attP* docker *y^−^w^−^M^eGFP,vas-int,dmRFP^ZH-2A;P{CaryP}attp40* and the balancer used in crosses *w^1118iso^/y+Y;Sco/SM6a;3^iso^* were used as controls for measuring DCV mortality and viral titre. Two replicates (A and B) for each of the two *pst* alleles were established from independent transformation events. Western blot with FLAG antibody were carried out using adult flies that were kept in 25°C to confirm the expression of FLAG-tagged *pst* alleles.

To assay the susceptibility of these lines to *pst*, vials were set up containing 10 females and 10 males of the transgenic lines and kept in 25°C. The parental flies were removed and the progeny collected. 15 vials containing 20 3-5 days old mated females of each line were inoculated with DCV (or Ringer’s solution as a control) as describe below, and their mortality was monitored for 19 days. Meanwhile, 15 additional vials of each line with 15 mated females were inoculated with DCV and maintained at 25°C. At day 2 post infection (day 6 at 18°C), total RNA of these flies were extracted and used to measure viral RNA levels by real-time PCR (see below).

### Measuring pst expression in DGRP lines

To study natural variation in gene expression, we measured *pst* expression in a panel of inbred fly lines from North America called the DGRP lines. We assayed 196 fly lines using one to seven biological replicates (a total of 654 RNA extractions). The flies were aged 6-9 and a mean of 15 flies was used for each RNA extraction. These RNA extractions had been generated as part of a different experiment and were infected with Nora virus (*pst* is not associated with susceptibility to Nora virus; R. Cogni, pers. comm. and expression of *pst* is not affected by Nora virus infection (Cordes, et al. 2013)). Primers and probes used are described below.

### Genotyping and naming of SNPs

DNA was extracted using either DNeasy Blood & Tissue Kit (Qiagen) according to manufacturer protocols or using a Chelex extraction that involved digesting fly tissues for 1 hr at 56°C with 5% w/v Chelex 100 ion exchange resin (Bio-Rad, Hercules, CA) in 200 ul of 33 mM dithiothreitol with 20 ug proteinase K (Jiggins and Tinsley 2005).

Diagnostic primers were designed to amplify *pst* allele carrying specific SNPs. The SNP of interest was put at the 3’ end of one primer and at least one mismatch next to the SNP was introduced (Table S2). In order to experimentally confirm the structural variants of *pst*, primers were designed to overlap the breakpoints of duplications and deletions (Table S2). PCR products were run on 1% w/v agarose gels.

SNPs were named according to their position in the *pst* gene. Numbering begins at the nucleotide encoding the start of the 5’ UTR, and includes intronic positions. The numbering of duplications and deletions refers to the size of the region affected in nucleotides. In the text we also report the genome coordinates of all variants.

### Drosophila C Virus

DCV stain C (Jousset, et al. 1972) was kindly provided by Luis Teixeira (Teixeira, et al. 2008)and was cultured in *Drosophila melanogaster* DL2 cells using the protocol describe in Longdon et al. (Longdon, et al. 2013). The Tissue Culture Infective Dose 50 (TCID50) was calculated by the Reed-Muench end-point method (Reed LJ 1938).

### Infection and resistance assay

Newly emerged flies were tipped into new food bottles. Two days later, mated females were infected with DCV by inoculating them with a needle dipped in DCV suspension as described in Longdon et al. (Longdon, et al. 2013) (TCID50=10^6^). Infected flies were kept on cornmeal food without live yeast on the surface. Numbers of infected flies that died were recorded every day and surviving flies were tipped onto new food every 3 days. Flies that died within 24hr were excluded from the analysis as it was assumed that they died from the injection process. Infected flies were collected on day 2 post infection for the measurement of viral titres.

### Quantitative RT-PCR

RNA was extracted using TRIzol (Invitrogen Corp, San Diego) in a chloroform-isopropanol extraction following the manufacturer’s instructions. RNA was used as template in qRT-PCR using QuantiTect Virus+ROX Vial Kit (^©^QIAGEN). Dual labelled probes and primers were ordered from Sigma-Aldrich^®^. The PCR primers and probes amplifying both the reference gene and the gene of interest were multiplexed in a single PCR reaction. DCV titre was measured using probe DCV_TM_Probe ([6FAM]CACAACCGCTTCCACATATCCTG [BHQ1]) and primers DCV_qPCR_599_F (5’ GACACTGCCTTTGATTAG 3’) and DCV_qPCR_733_R (5’ CCCTCTGGGAACTAAATG 3’). The amount of virus was standardised to a reference gene *RPL32* using probe Dmel_RpL32_TM_Probe ([HEX]ACAACAGAGTGCGTCGCCGCTTCAAGG[BHQ1]) and primers: Dmel_RpL32_F (5’ TGCTAAGCTGTCGCACAAATGG 3’) and Dmel_RpL32_R (5’ TGCGCTTGTTCGATCCGTAAC 3’). Expression of *pst* was measured using dual labelled probe Pst_PR ([Cy5]CAGCACACCATTGGCAACTC [BHQ3]) and primers Pst_FW (5’ CCGTCTTTTGCTTTCAATA 3’) and Pst_RV (5’ CCCAACTGACTGTGAATA 3’). The amount of *pst* expression was standardised to a reference gene *Ef1alpha100E* using the ΔΔCt (critical threshold) method (see below). Expression of *Ef1alpha100E* was measured using probe ([FAM] CATCGGAACCGTACCAGTAGGT [BHQ2]), primers Ef1alpha100E_FW (5’ ACGTCTACAAGATCGGAG 3’) and Ef1alpha100E_RV (5’ CAGACTTTACTTCGGTGAC 3’). Subsequent to the experiment we realised there was a SNP segregating in the sequence to which the probe Pst_PR annealed, so the effect of this was corrected for by estimating the effect of this by linear regression and correcting the ΔCt; values for its effect. This procedure did not qualitatively affect the conclusions. The estimation of gene expression or viral titre assumed that that the PCR reactions were 100% efficient. To check whether this assumption is realistic we used a dilution series to calculate the PCR efficiency. Three technical replicates of each PCR were performed and the mean of these used in subsequent analyses. All the PCR efficiencies were between 97%-103%.

### Statistical analysis of survival data and viral titres

R version 3.2.1 (Team 2008) was used for statistical analyses. In the experiments using flies overexpressing *pst* and or flies transformed with a modified BAC clone we recorded the lifespan of individual flies. This data was analysed with a Cox proportional hazard mixed model, fitted using the R package “coxme”. The genotype of the fly line was treated as a fixed effect. The random effects were the vial in which a fly was kept which was nested in the replicate fly line (where the same fly genotype had been generated twice by independent transformation events). Flies alive at the end of the experiment were censored.

For each fly line in which we measured viral titres by quantitative RT-PCR, we first calculated ΔCt as the difference between the cycle thresholds of the gene of interest and the endogenous controls (*actin 5C* or *Ef1alpha100E*). We used the mean values of technical replicates. To assess whether these differences were statistically significant, we fitted a general linear mixed model using the *lme* function in R. We used the mean Δ*Ct* across all biological replicates as a response variable. The genotype of the fly line was treated as a fixed effect and the day that the flies were injected as a random effect.

### Identifying structural variants of pst

We identified structural variants by looking at the sequence data of the DGRP genomes (Mackay, et al. 2012). Structural variants were detected when two halves of the same sequence read or read-pair map to different positions or orientations within the reference genome. We analysed 205 Freeze 2 BAM files of the DGRP lines (Mackay, et al. 2012) using Pindel_0.2.0 (Ye, et al. 2009) to identify the breakpoints of structural variants among the lines (deletions, tandem duplications, large and small insertions). In 192 of the 205 lines we confirmed the structural variants by carrying out PCR using primers either overlapping breakpoints or flanking them (Table S2). We repeated this twice for the small number of lines that showed conflicting results with the Pindel analysis. We also Sanger sequenced the breakpoints in a subset of lines to confirm the predictions from the short read analysis.

Duplications and deletions can also be detected by changes in sequence coverage. A script written in Python was used to calculate the coverage number for each base pair in the region 3L:7338816-3L:7366778 (BDGP 5).

### Identifying multiple alleles of pst with different effects on DCV susceptibility

We have previously measured survival after DCV infection in a panel of inbred fly lines called the *Drosophila* Synthetic Population Resource (DSPR) Panel B (King, et al. 2012a; King, et al. 2012b). These lines were constructed by allowing 8 inbred founder lines with complete genome sequences to interbreed for 50 generations, and then constructing inbred lines (RILs) whose genomes were a fine-scale mosaic of these founders. We infected 619 RILs in Panel B with DCV and monitored the mortality of 14091 flies post infection, which allowed us to identify *pst* as a major-effect gene defending flies against DCV infection (Cogni, et al. 2016).

In this study, we reanalysed this dataset to test whether there were more than two alleles of *pst*. To identify different alleles of *pst*, we used a Hidden Markov Model (HMM) (King, et al. 2012a) to determine which of the 8 founder lines the *pst* allele had been inherited from. We assigned RILs to one of the founders when position 3L: 7350000 (the location of *pst*) could be assigned to that parent with ≥ 95% confidence. We analysed this data with a one-way ANOVA, with the mean survival time of each vial RIL as the response variable, and founder allele as a fixed effect. We then performed a Tukey’s honest significant difference test to assign the founders into allelic classes with differing levels of resistance.

### Identifying cis-regulatory polymorphisms in pst

To look for cis-regulatory polymorphisms that cause variation in *pst* expression, we used a set of microarray data of female head tissue in the DSPR (King, et al. 2014). The mean normalised expression of 3 *pst* probes that did not contain any SNPs segregating in the panel (FBtr0273398P00800, FBtr0273398P01433, FBtr0273398P01911) were used. The QTL analysis was performed using the R package DSPRqtl (http://FlyRILs.org/Tools/Tutorial) (King, et al. 2012b) following Cogni *et al* (Cogni, et al. 2016).

### Association between pst expression and DCV resistance

To test whether the structural variants were associated with *pst* expression or susceptibility to DCV, we genotyped 192 DGRP lines (which flies were available then) for structural variants by PCR (primers listed in Table S2). These variants were then combined with sequence data from the DGRP lines (Freeze 2). We have previously measured the survival of these fly lines after DCV infection (Magwire, et al. 2012). The mean *pst* expression level was measured in 196 DGRP lines (see above). We then tested for associations between the SNPs in the region of 3L: 7311903-7381508 (BDGP5) and the mean of *pst* expression of each DGRP line using a linear model.

To estimate the genetic correlation between *pst* expression and survival after DCV infection in the DGRP lines we used a bivariate general linear mixed model. The mean survival time of flies post DCV infection was calculated for each vial assayed. *Pst* expression was measured on whole vials of flies. *pst* expression and survival as Gaussian response variables in the model:

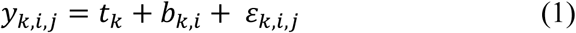

Where *y_i,j,k_* is the observed trait *k (pst* expression level or mean survival time) of flies from line *i* in vial *j. t_k_* is a fixed effect representing the mean expression level (Δ*Ct*) or survival time. *b_ki_* are the random effects, which are assumed to be multivariate normal with a zero mean. For the random effects we estimated a 2x2 covariance matrix describing the genetic (between-line) variances of *pst* expression and survival, and the covariance between these traits. The genetic correlation was calculated from these parameters. *E_k,i_,_j_* is the residual error, with separate residual variances estimated for the two traits. The parameters of the models were estimated using the R library MCMCglmm (D. 2010), which uses Bayesian Markov Chain Monte Carlo (MCMC) techniques. Each model was run for 1.3 million steps with a burn-in of 300,000 and a thinning interval of 100. Credible intervals on all parameters (variances, correlations etc) were calculated from highest posterior density intervals. The analysis was repeated including SNP A2469G as a fixed effect to control for any confounding effects of this variant being in linkage disequilibrium with *cis*-regulatory polymorphisms (assuming this SNP is not itself a *cis* regulatory polymorphism).

### Test for natural selection on pst

To investigate the frequency of resistance allele of A2469G in populations worldwide, we looked at publically available genome resequencing datasets of the Global Diversity Lines (Grenier, et al. 2015), North American population (DGRP (Mackay, et al. 2012)) and Zambian population (DPGP (Pool, et al. 2012)). We also collected 341 iso-female *D. melanogaster* from Accra, Ghana and genotyped a pool of flies from these lines for SNP A2469G by PCR as described above.

To test for signature of natural selection on *pst*, we analysed the sequence around *pst* from publically available genome sequences of *Drosophila*. These sequences were either from inbred lines or haploid genomes, so the data was phased as haplotypes. We analysed data from two populations of *D. melanogaster* with large sample sizes: a North American population (DGRP, 205 lines) and a Zambian population (DPGP3, 196 lines). The variant calls from these lines in VCF file format of freeze2 DGRP was downloaded from BCM-HGSC website (Mackay, et al. 2012). Because duplication and rearrangement of *pst* is very common in *D. melanogaster* (Figure 3), in the DGRP lines we Sanger sequenced *pst* from 35 lines of variant 3 and 28 lines of variant 4 so we only analysed data from the complete copy of the gene. These sequences were combined with 105 DGRP lines without rearrangement, resulting in a total of 165 DGRP lines with *pst* sequences. This was not possible for the data from Zambia as the original lines are not available. Here, consensus sequences of 196 *D. melanogaster* samples were downloaded from http://www.johnpool.net/genomes.html/. About 20kb sequence around *pst* (3L: 7340375-7363363) were pulled out from all lines using the scripts “breaker.pl” and “dataslice.pl” written by the authors, returning FastA files. Then FastA file was converted into VCF file by PGDSpider (Lischer and Excoffier 2012).

To examine how allele frequencies differ between populations, *F_ST_* was calculated on a per-site basis for a North American population (DGRP) and a Zambian population (DPGP3) by VCFtools (Danecek, et al. 2011). To detect linkage disequilibrium (LD) around SNP A2469G, we estimated LD between all pairs of SNPs in 20kb region around it. R package “genetics” and “LDheatmap” (Shin J-H 2006) were used to calculate and plot LD an heat map. We then applied Long-Range Haplotype test (Sabeti, et al. 2005; Zeng, et al. 2007) to examine the extent haplotype homozygosity (EHH) around SNP A2469G in comparison to other haplotypes of similar frequency in the 200kb region (3L: 7250375-7253363). R package “rehh” was used in the analysis (Gautier and Vitalis 2012).

We finally applied a McDonald and Kreitman (MK) test to detect positive selection on the amino acid level (McDonald and Kreitman 1991). Using the *D. yakuba* sequence as an outgroup, substitutions were polarised along the lineage leading from the common ancestor of *D. melanogaster* and *D. simulans* to *D. melanogaster*. A standard MK test was carried out using McDonald and Kreitman Test (MKT) software (Egea, et al. 2008). We excluded polymorphic sites with a frequency less than 10% to reducse the number of deleterious amino acid polymorphisms in the dataset. Polarized 2 × 2 contingency tables were used to calculate α, which is an estimate of the proportion of amino acid substitutions fixed by selection (Smith and Eyre-Walker 2002). Statistical significance of the 2 × 2 contingency tables was determined using a χ^2^ test.

Nucleotide diversity was calculated using DnaSP v5 (Librado and Rozas 2009) for the 20 kb region described above in 165 DGRP lines and 196 DPGP lines.

## Results

### The pst locus has multiple alleles affecting DCV resistance

In a previous association study we found 6 SNPs in *pst* that were strongly associated with DCV resistance (Magwire, et al. 2012). All of these are in linkage disequilibrium (LD) with each other, so it was not possible to identify the causative variant from this data. Intriguingly however, no single SNP could explain all the effects of *pst* on DCV susceptibility, suggesting that multiple alleles of this gene with different susceptibilities might be segregating in populations. To investigate this further, we reanalysed a second dataset where we had infected 13,919 flies from the *Drosophila* Synthetic Population Resource (DSPR, panel B, 619 Recombinant inbred lines founded by 8 lines representing a worldwide sample (King, et al. 2012b)) with DCV and shown that resistance was largely controlled by *pst* (Cogni, et al. 2016). These data allow us to estimate the effect that each of the 7 different founder haplotypes of *pst* segregating among these lines has on DCV susceptibility (one of the eight founders, BB5 was removed from analysis because it is represented by less than 10 lines and was not able to be assigned to any group). The 7 founder haplotypes fall into 4 groups with significantly different resistance levels (Table 1). Flies in resistant 1 (resist1) group survived an average of 9.6 days post infection while flies in resistant 2 (resist2) group survived an average of 11 days post infection. Flies in susceptible 1 (susc1) group survived an average of 6.1 days post infection while flies in susceptible 2 (susc2) group survived an average of 7.1 days post infection (Table 1). Therefore, in a sample of seven copies of this gene, there are four functionally distinct alleles of *pst* affecting DCV resistance.

**Table 1.**
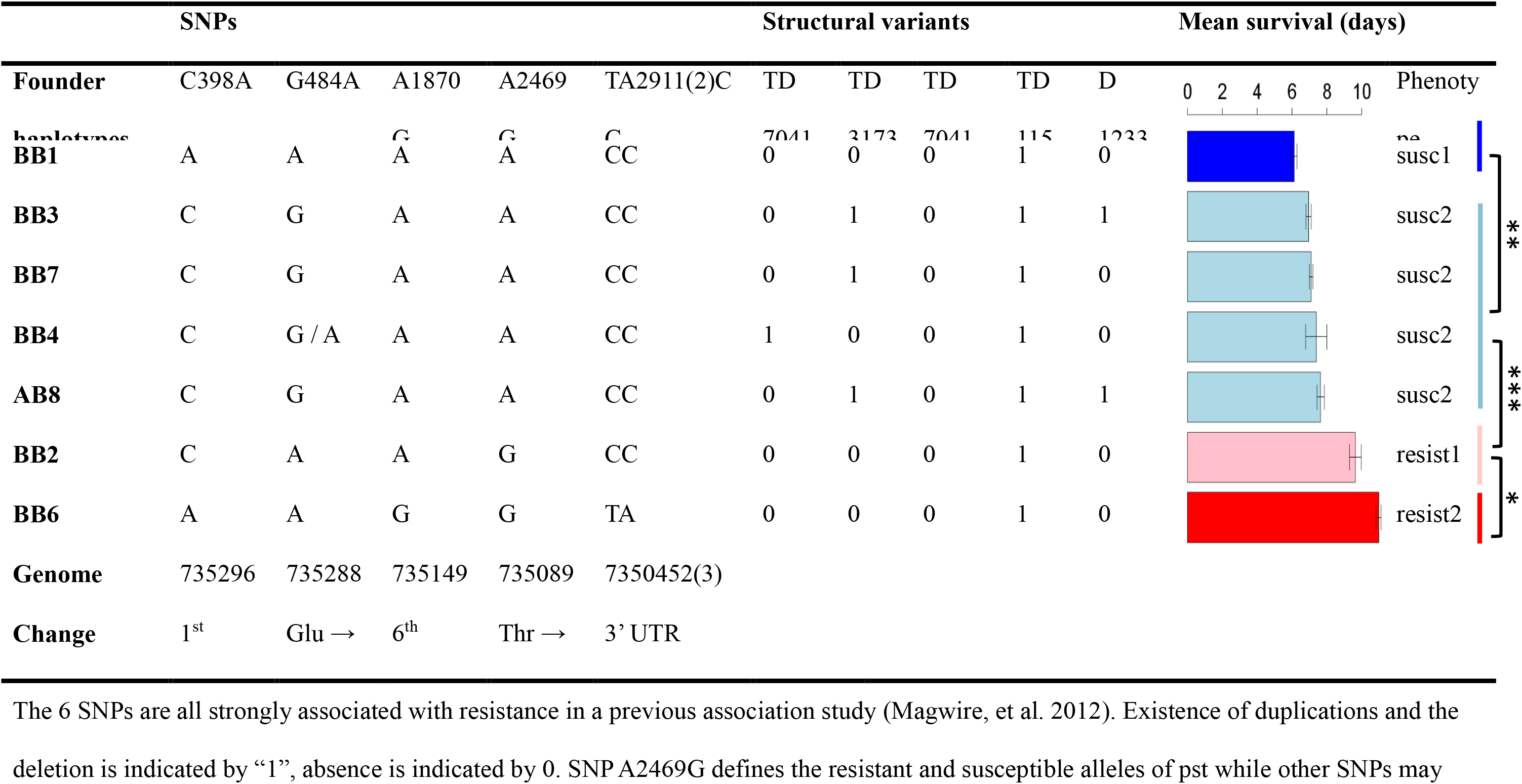
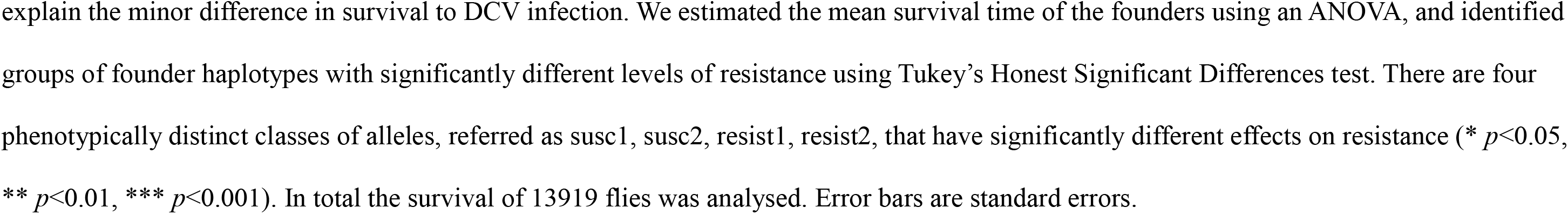
Candidate SNPs and structural variants in the 7 founder haplotypes segregating in the DSPR panel.

### The amino acid substitution A2469G can explain resistance in two different genetic mapping experiments

We examined the 6 *pst* SNPs previously found associated with resistance in our genome-wide association study in DGRP lines (*p*<10^-12^) and asked which of them explain the four levels of resistance we observed in the DSPR founders. Only A2469G, which is a non-synonymous change (Thr/Ala, 3L: 7350895, BDGP5), can explain the large difference between the two resistant and the two susceptible classes of alleles (Table 1). This change is also the most significant SNP in the association study using the DGRP lines (Magwire, et al. 2012) and is a separate study that had selected populations for DCV resistance and then sequenced their genomes (Martins, et al. 2014). This threonine to alanine change is a radical substitution between a polar and a nonpolar amino acid, and alanine is associated with increased resistance in both the association study and this QTL analysis. Two closely related species *Drosophila simulans Drosophila yakuba* both have a threonine at this position, indicating that the susceptible allele was the ancestral state. While this analysis strongly implicates A2469G in resistance, it does not preclude a role for the other 5 variants associated with resistance. For example, SNP C398A differs between the susc1 and susc2 alleles, while SNPs TA2911(2)CC, A1870G and C398A all differ between resist1 and resist2.

### Modifying SNP A2469G in transgenic flies confirms that it alters resistance to DCV

To experimentally confirm the SNP(s) causing flies to be resistant to DCV, we generated five transgenic lines where we modified each of the six SNPs associated with resistance (SNP T2911C and A2912C, which are in complete LD, were modified together: TA2911(2)CC). To do this we edited a BAC clone (*CHORI-322-21P14*, 20.064kb) of the region in *E. coli*. The BAC originally contains the allele associated with increased resistance for all the five *pst* variants, and we individually changed these to the susceptible variant. We inserted the five BACs into the same genomic position in a fly line to generate five transgenic lines. We crossed these transgenic flies to a balanced *pst* hypomorphic mutant *y^1^w^67c23^;If/Cyo;P^GSV1^GS3006/TM3,Sb^1^Ser^1^*, which has a transposable element inserted in the 5’ UTR of the *pst* gene. The transgenic alleles did not complement the lethal effect of this mutation upstream of *pst*, so we infected flies that were homozygous for the transgenic *pst* allele on chromosome 2 and had one hypomorphic mutant allele over a balancer chromosome on chromosome 3 *(pst* hypomorphic mutant allele and the balancer carries the susceptible form “A” for SNP A2469G).

The two independent fly lines carrying a “A” for SNP A2469G, which were generated through independent transformation events, died significantly faster after DCV infection compared to all the other transgenic lines that had a “G” at this position (Figure 1). There were no significant differences among the other 4 genotypes. Among the flies that were mock infected with Ringer’s solution there were no significant differences among lines (although flies carrying a susceptible “A” at A2469G survived longest, which reinforces the result that the high mortality of these flies when DCV infected is being caused by *pst*). In summary, both genetic mapping approaches and experimentally modifying the gene demonstrate that the SNP A2469G is causing flies to be resistant to DCV.

**Figure 1.**
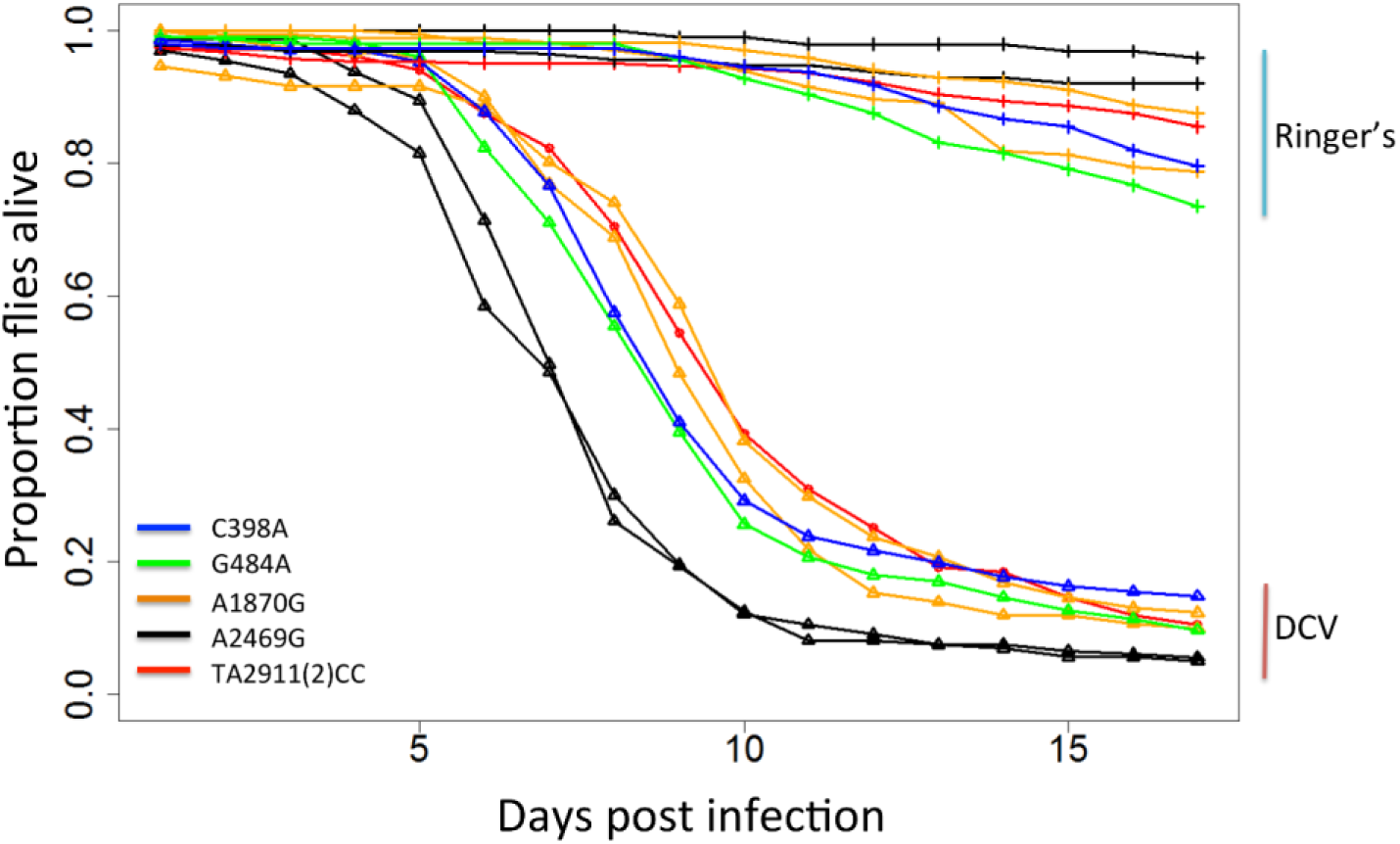
Susceptibility to DCV in transgenic flies carrying different alleles of *pst* SNPs. Lines with triangles are flies infected with DCV while lines with crosses are flies injected with Ringer’s solution as a control. SNP A2469G and SNP A1870G have two biological replicates, which were generated through independent transformation events. By fitting a Cox proportional hazard mixed model, we found that A2469G is significantly different from all the other SNPs (*P*<0.007). There were no significant differences among the other 4 SNPs (*P*>0.36). In total, 157 vials containing 3010 females were infected with DCV and their mortality were recorded daily for 17 days.

### Overexpressing both the resistant and susceptible alleles of pst protects flies against DCV infection

Resistance could evolve by altering host factors that are beneficial to the virus or by increasing the efficacy of existing antiviral defences. To distinguish these hypotheses, we generated fly lines that overexpress either the resistant or the susceptible allele of *pst* (these constructs encode a protein that only differs at the site affected by SNP A2469G). The two FLAG-tagged constructs were inserted at the same position of the fly genome using *phiC31* integrase, and we checked that the full length protein (approximately 77kDa) was being expressed using a western blot targeting the FLAG tag. Two replicates of these lines were generated and these flies were then infected with DCV. We found that overexpressing both the susceptible and the resistant alleles of *pst* led to significant reductions in viral titres at two days post-infection (Figure 2A; General linear model: *pst*_A_: |z|=3.3, *P*=0.003, *pst*_G_: |z|=4.83, *P*<0.001). There is no significant difference in viral titre between flies overexpressing the resistant *pst* allele “G” and flies overexpressing the susceptible *pst* allele “A”, although the trend is in the expected direction (Figure 2A; |z|=1.4, *P*=0.35). Next, we examined survival. Overexpressing *pst*, no matter which allele, substantially increased survival after DCV infection (Cox proportional hazard mixed models; *pst_A_:* |z|=12.32, *P*<1e^-5^, *pst_G_*: |z|=11.83, P<1e^-5^) (Figure 2B). Again, we were not able to detect any difference in mortality between flies overexpressing the two different alleles of *pst* (|z|=0.53, *P*=0.86). This result should be interpreted with caution, as any differences in resistance between the two alleles may be obscured by intrinsically lower survival of the flies overexpressing the resistant allele (Figure 2B, Ringers control). This difference in the survival of mock-infected flies overexpressing the different alleles could not be replicated when new transgenic flies were generated in a different genetic background and assayed without pricking, suggesting that it is not a toxic effect of the resistant allele (Figure S1). In summary, overexpressing either *pst* allele substantially increased resistance, with the resistant allele causing a slightly greater reduction in viral titre.

**Figure 2.**
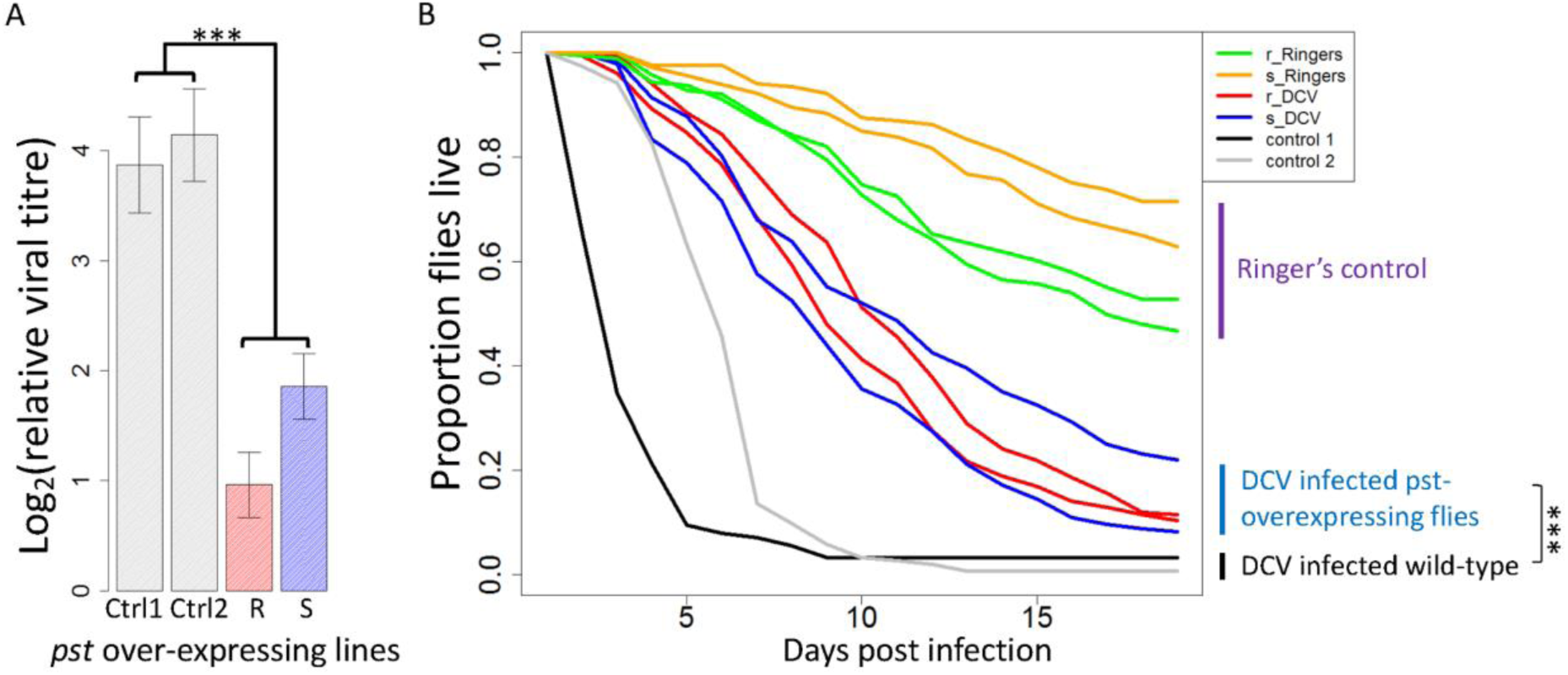
The effect of over-expressing pst carrying the susceptible (s) and resistant (r) alleles of SNP A2469G on survival and viral titre. (A) DCV titre relative to *Act5C* in flies 2 days post infection. Bars are the means of 28 vials each containing 15 flies. Error bars are standard errors. (B) The proportion of flies alive after infection with DCV or mock infection with Ringer’s solution. The survival curves are the mean of ~15 vials of flies, with a mean of 18 flies in each vial. Flies were kept at 25°C. Control 1 (Ctrl1) were docker flies which the BAC constructs inserted into *(y^−^w^−^Me^GFP,vas-int,dmRFP^ZH-2A;P{CaryP}attp40*), and Control 2 (Ctrl2) were flies used in the crosses to select successful transformants *(w^1118iso^/y^+^Y;Sco/SM6a;3^iso^)*. The experiments used two independent transformants of each construct (A and B). ^*^stands for *P*<0.05, ^***^ stands for *P*<<0.001.

### The pst locus contains complex structural polymorphisms

The analyses above only considered SNPs, but other types of genetic variation could cause flies to be resistant. We therefore investigated the existence of structural variation in a panel of 205 inbred fly lines from North America whose genomes had been sequenced (DGRP) (Mackay, et al. 2012). The existence of structural variation had been suggested by the PCR amplification of a truncated copy of *pst* in certain flies and cell lines. We identified the breakpoints of structural variants from published paired-end short read sequencing data (using Pindel_0.2.0) (Ye, et al. 2009). Excluding small indels, this approach revealed 5 variants that were shared by more than 2 lines and supported by at least 4 raw sequencing reads (Figure 3A; the region investigated, 3L: 7,346,678-3L: 7,357,466, DPGP 5, includes *pst* and the two flanking genes *CTCF, Sec63*). In 192 of the 205 lines we confirmed the structural variants by carrying out PCR with diagnostic primers and Sanger sequencing. As a final confirmation we checked that the duplicated regions had increased sequence depth (Figure 3B).

**Figure 3.**
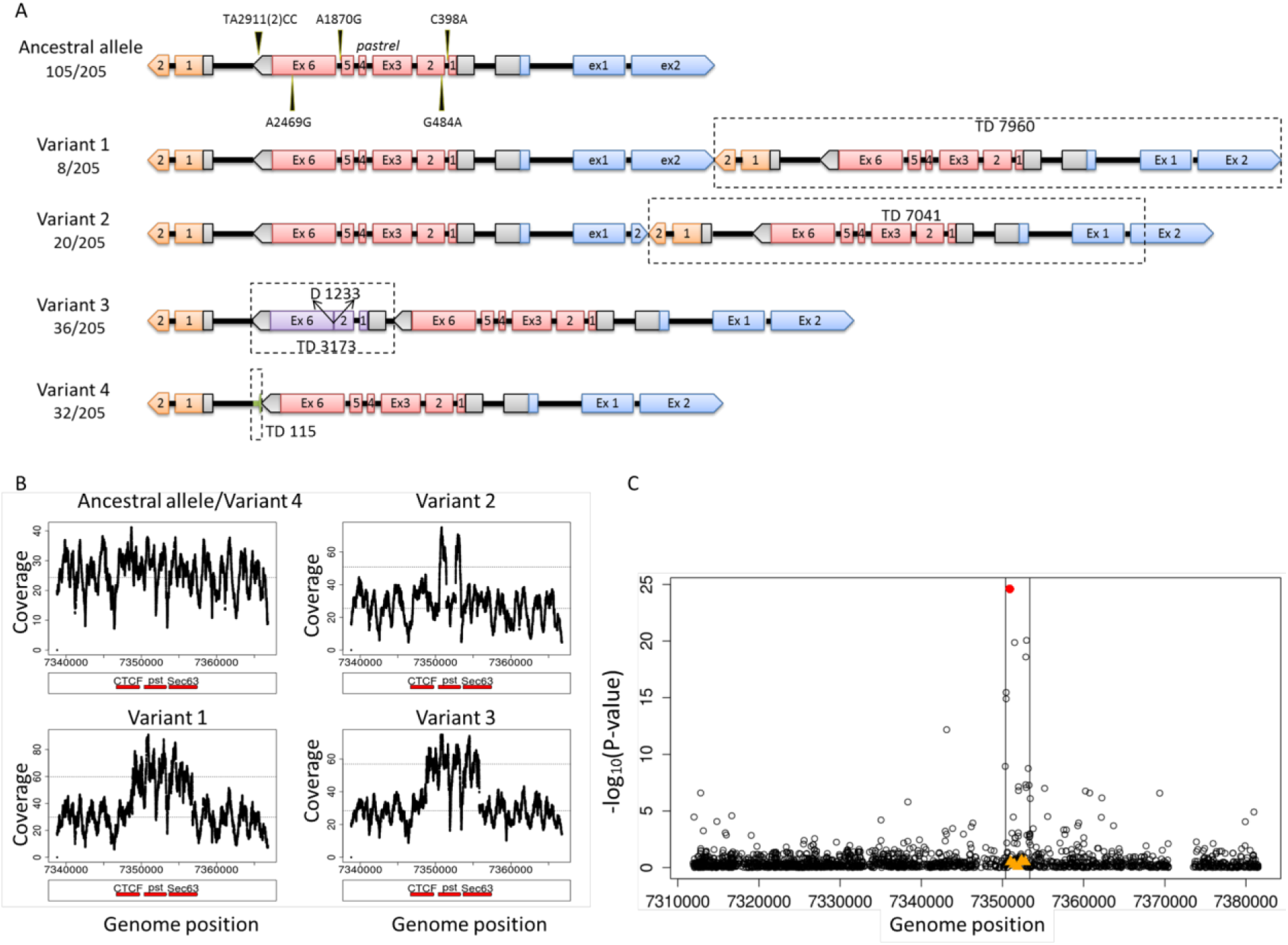
Five structural variants of *pastrel*. (A) Cartoon of *pst* variants, with alleles’ size scaled to gene length. Pink boxes represent complete copy of *pst* gene; orange boxes represent coding sequence of gene *CTCF* located at 3’ end of *pst*; blue boxes represent coding sequence of gene *Sec63*, located at 5’ end of pst; grey boxes are UTRs; purple boxes are truncated copy of *pst* gene. Allele frequencies in DGRP are shown below the variant name. Variant 2 differs from variant 1 in that variant 2 has a shorter duplication of CTCF exon2. (B) Mean sequencing coverage plots of the region 3L: 7,338,816 (1kb upstream of the start of TD7960) - 7,366,778 (1 kb downstream of the end of TD7960) for ancestral allele of *pst* and four structural variants. Red bars stand for *pst* and two neighbour genes *CTCF* and *Sec63*. Variant 4 has a very short duplication 115 bp so shows very similar coverage plot as the ancestral allele. Sequence data is from the original DGRP genome sequencing project **(Mackay, et al. 2012)**. (C) Association between survival after DCV infection and *pst* SNPs and structural variants. -Log_10_ (p-value) of the association between SNPs in the region of 3L: 7311903 - 3L: 7381508 (BDGP 5) and survival is plotted against genome positions of the SNPs. SNPs are showed as empty circles; SNP A2469G is in red. Structural variants of *pst* are showed in orange triangles.

The five major structural variants and their frequencies in the DGRP lines are summarised in Figure 3A. Just over half the lines had the ancestral state that is found in the reference genome with one complete copy of *pst* (Figure 3A; ancestral allele). 8 out of 205 lines have a 7960bp duplication (3L: 7348816-7356777, variant 1) containing a complete copy of *pst* and some sequences from two adjacent genes (*CTCF, Sec63*). 20 lines have a 7041bp duplication (includes *pst* and partial sequences from neighbour genes, 3L: 7348778-7355820, variant 2). 36 lines have a duplicated copy of *pst* (3L: 7350246-7353420) with a 1233bp deletion in the middle (variant 3). 32 lines have a duplication of 115bp (3L: 7350263-7350379) at the 3’ end of *pst* (variant 4). There are another 4 lines containing structural variants each represented by less than 3 lines that are not shown in Figure 3A.

We tested whether these structural variants affect survival of the DGRP lines after DCV infection and found that none of the structural variants is associated with survival post DCV infection (F_1_,_165_<1.97, *P*>0.32; Figure 3C). This non-significant result may due to that we lack the power to detect their effects. For example one of the structural variants that has a complete copy of *pst* was only represented by as few as 8 lines. Another possible explanation is that transcripts produced by these duplicates are non-functional.

### There is cis-acting genetic variation that alters the expression of pst

Given that altering the expression of *pst* experimentally alters resistance to DCV, it is possible that natural variation in gene expression affects susceptibility to the virus. We investigated this using published microarray data from F1 individuals from crosses between two panel of recombinant fly lines derived from 15 founder lines from around the world (crosses between DSPR panel A females and panel B males) (King, et al. 2014). To map regions of the genome affecting *pst* expression, we used the mean normalised expression of 3 *pst* probes (FBtr0273398P00800, FBtr0273398P01433, FBtr0273398P01911) that did not contain any SNPs. We found that there was a major QTL controlling *pst* expression at 3L: 7350000 (LOD=35.04), which is very close to the location of *pst* (Figure 4A). Therefore, there is genetic variation in *pst* expression and this is controlled by *cis*-acting genetic variants close to *pst* rather than variation elsewhere in the genome acting in *trans*.

**Figure 4.**
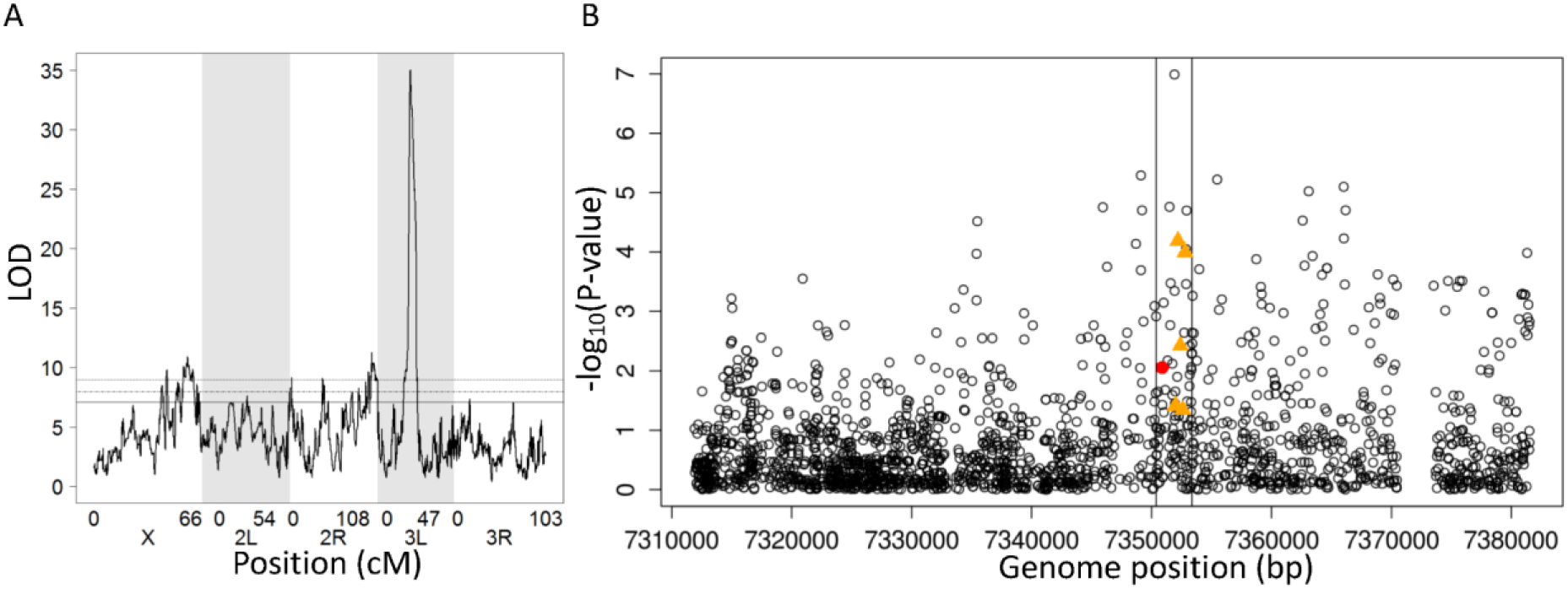
Cis-regulatory variation in pst expression. (A) Map of QTL associated with pst expression in female head of DSPR crosses. A single peak at position 3L: 7350000 was found (LOD=35). The horizontal line is the genome-wide significance threshold obtained by permutation (p<0.05, LOD=7.12). Expression data is from published microarray analysis (King, et al. 2014). (B) Association between pst expression and its SNPs and structural variants in DGRP lines. Gene expression was measured by quantitative RT-PCR on 654 biological replicates of 196 fly lines. -Log_10_ (*p*-value) for the association between SNPs and expression in the region of 3L: 7311903 - 3L: 7381508 (BDGP 5) is plotted against genome positions of the SNPs. SNPs are showed as empty circles; SNP A2469G in red. Structural variants of *pst* are showed in orange triangles.

To investigate which genetic variants might be affecting *pst* expression, we measured expression across 198 DGRP lines using quantitative RT-PCR. Using this data we looked for associations with the 5 structural variants and SNPs in the region surrounding *pst* (Figure 4B). We found *pst* expression was most significantly associated with a SNP in an intron of *pst* at position A1455T (3L: 7351909, *F_2,155_*=17.89, *P*=1.02e^-7^). However, several of the structural variants were also associated with *pst* expression (Figure 4B). Tandem duplication TD3173 and TD115 were in linkage disequilibrium with SNP A1455T (Fisher’s Exact Test: *P*=0.002 and *P*=0.001), but they remain significantly associated with *pst* expression after accounting for SNP A1455T by including it as a covariate in the model (TD3173: *F*_1,156_=14.07, *P*=0.0002; TD115: *F*_1,156_=7.7*, P*=0.006; Figure 4B). TD3173 is in strong linkage disequilibrium with D1233 (Fisher’s Exact Test, *P*<2.2e^-16^). Therefore, multiple *cis* regulatory variants affect *pst* expression, and these may include structural variants.

### The expression of pst is correlated with DCV resistance

Across 198 DGRP lines we found that natural variation in *pst* expression was correlated with survival after DCV infection (Figure 5; Genetic correlation: *r_g_*=0.32, 95% CI=0.17 to 0.45). This is consistent with previous results that DCV resistance changes when *pst* is knocked down by RNAi (Magwire, et al. 2012) or over-expressed using transgenic techniques (Figure 2). SNP A2469G is a non-synonymous SNP that was found to affect survival after DCV infection, which means it is unlikely to have an effect on gene expression. However, if it is in linkage disequilibrium with a *cis* regulatory variant this could create spurious associations between gene expression and resistance. To control for this we estimated the correlation after accounting for the effect of SNP A2469G by including it as a covariate in the model, and found the correlation between *pst* expression and survival after DCV infection remains significant (Genetic correlation: *r_g_*=0.25, 95% CI=0.11 to 0.41). These results indicate that *cis*-regulatory variation that alters *pst* expression and affects resistance to DCV.

**Figure 5.**
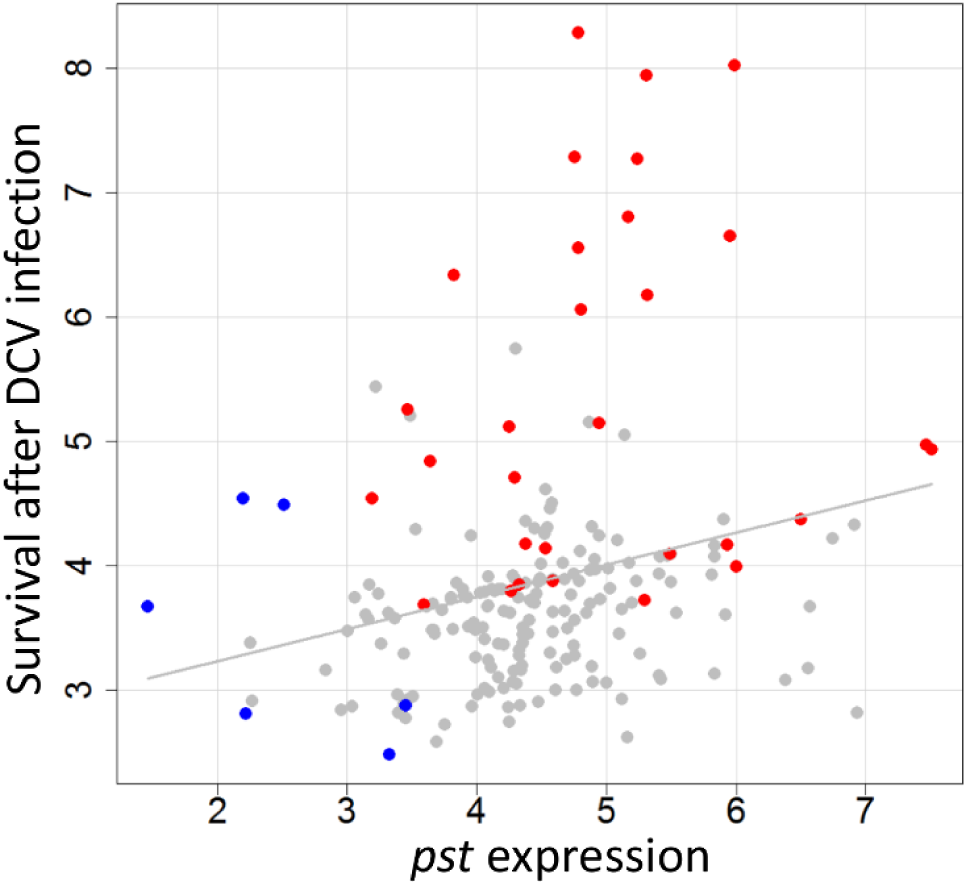
Correlation between *pst* expression and survival after DCV infection in DGRPs. Grey line is fitted by linear regression line and is shown for illustrative purposes only. Each point is the estimated phenotype of a single DGRP line (marginal posterior modes of the random effects in model equation 1). Red dots represent lines contain resistant allele “G” for SNP A2469G and blue dots represent lines contain “T” for SNP A1455T. Gene expression was measured by quantitative RT-PCR on 654 biological replicates of 196 fly lines. Survival after DCV infection was estimated from 730 vials of flies, with the data from Magwire et al (Magwire, et al. 2012).

### There is no evidence of spatially varying selection acting on the resistant allele of pst A2469G

Having identified the genetic variant that is responsible for most of the genetic variation in DCV resistance in *D. melanogaster*, we are well-placed to characterise how natural selection has acted on this variant. It is common to find that the prevalence of viruses in *Drosophila* varies geographically (Carpenter, et al. 2012; Webster, et al. 2015), and this is expected to result in spatially varying selection pressure for resistance. However, there is little variation in the frequency of the resistant allele between populations. The resistant allele of A2469G is at a low frequency in populations worldwide—7.7% in Zambia (197 DPGP3 lines (Pool, et al. 2012)), 16% in North America (205 DGRP lines (Mackay, et al. 2012)), 10% in Beijing (15 GDL lines (Grenier, et al. 2015)), 5% in the Netherlands (19 GDL), 33% in Tasmania (18 GDL) and 10% in Ghana (341 lines collected and genotyped in this study). Among the populations with genome sequence data, only Zambian and North American populations have large sample sizes (196 lines and 205 lines separately), so the following analysis were carried out on these two datasets.

To compare the geographical variation in allele frequency at A2469G to other SNPs in the region, we calculated *F_st_* (a measure of differences in allele frequency) between North America and Zambia. It is clear that A2469G (red star in Figure 6A) is not a significant outlier relative to the other 2641 SNPs analysed in the region 100kbp either side (SNPs that have a minor allele frequency below 5% were filtered out), indicating there is no evidence of population-specific selective pressure on SNP A2469G.

**Figure 6.**
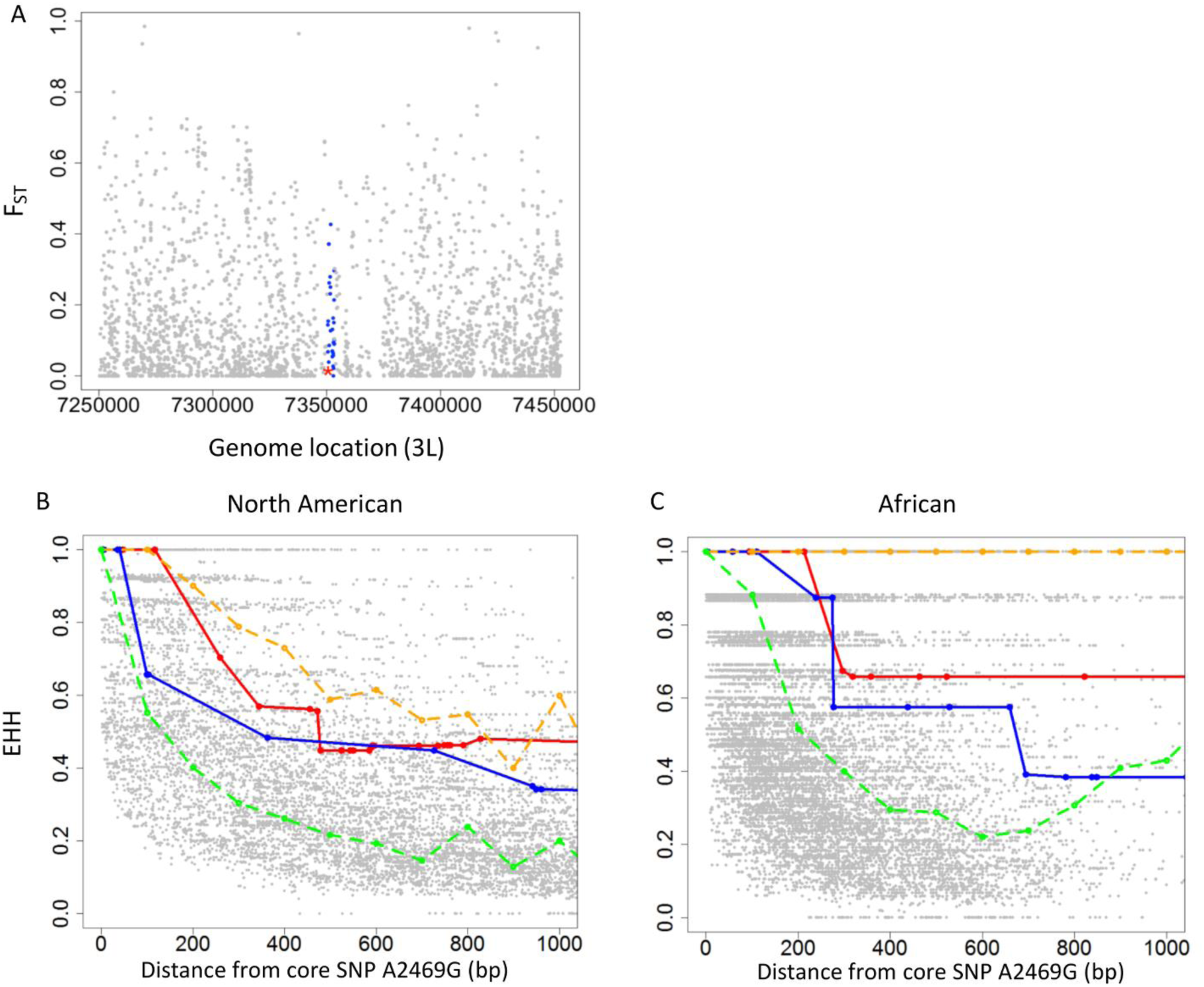
Population genetic evidence of natural selection acting on the amino acid polymorphism A2469G in Pastrel that confers resistance to DCV. (A) *F_ST_* of all SNPs within 200kb region around *pst*. Blue dots are SNPs in *pst*, and the red star is A2469G. *F_ST_* was calculated between Zambia and North America using published genome sequences (see text). Panels B and C show the breakdown of extended haplotype heterozygosity (EHH) over distance between the derived (resistant) allele of the core SNP A2469G and SNPs within the distance of 1000 bases from the mutation. Red line and blue line are EHH breakdown of upstream and downstream of SNP A2469G respectively. The grey points are a null distribution generated by calculating the EHH using other SNPs that are a similar frequency in the region as the core. The orange dash line indicates top 5% EHH value of this null distribution while green dash line indicates median EHH.

### The resistant allele of pst is old and shows no evidence of recent changes in frequency driven by natural selection

When natural selection causes an unusual rapid rise in allele frequency, there is little time for recombination to break down the haplotype carrying the selected mutation. This results in unusual long-range haplotypes and elevated linkage disequilibrium (LD) around the variant given its population frequency. As we know the site that is likely to be a target of selection, this is a powerful way to detect the effects of selection on DCV resistance. We first measured the LD between SNP A2469G and SNPs in a 10 kb region upstream and downstream of it. In both Africa and North America we found very little LD between SNP A2469G and surrounding SNPs (Figure S2 A and B).

When the variant under selection is known, the most powerful test for such effects is the EHH test (extended haplotype homozygosity) (Sabeti, et al. 2005; Zeng, et al. 2007). We calculated the EHH using the resistant (derived) allele of SNP A2469G as a core, and compared this to a null distribution generated from other SNPs of similar frequency that were nearby in the genome (Figure 6 B and C). In both populations, although the EHH around the resistant allele of A2469G is above the median, it is below the top 5%. Therefore, there is no evidence of positive selection on the resistant allele of A2469G generating extended LD around this variant. We also calculated the EHH for the susceptible allele of SNP A2469G as a core, and found no extended LD around this variant (Figure S3).

Positive and balancing selection can also affect the nucleotide diversity (π). In a 20kb region around *pst* in both North American and African populations we did not observe elevated nucleotide diversity compared to π value of whole genome (Figure S4 A and B). We also calculated π among chromosomes carrying the resistant or the susceptible allele of A2469G, and did not find altered patterns of diversity around *pst* (Figure S4 C and D).

It is common to find components of the immune system where natural selection has driven rapid evolution of the protein sequence, which is normally interpreted as being caused by selection by pathogens (Obbard, et al. 2009). To test whether this was the case for *pst* we tested whether other amino acid variants had been fixed in *pst* using the McDonald-Kreitman Test (McDonald and Kreitman 1991). *Drosophila yakuba* and *Drosophila simulans* sequences were used to infer the sequence of the most recent common ancestor of *D. simulans* and *D. melanogaster*. Analysing polymorphisms from 165 lines from the DGRP panel and divergence from the most recent common ancestor of *D. simulans* and *D. melanogaster*, we found no signature of positive selection (low frequency variants excluded; Synonymous Polymorphism=7, Synonymous Divergence=13.23, Non-synonymous Polymorphism=13, Non-synonymous Divergence=32.46, *α*=0.76, *χ*^2^=0.242, P=0.625). Therefore, there is no evidence of positive selection on the amino acid sequence of Pastrel over the last ~3 million years.

## Discussion

It has been argued that susceptibility to infectious disease may frequently have a simpler genetic basis than many other quantitative traits because natural selection drives major-effect resistance alleles up in frequency in populations (Hill 2012; Magwire, et al. 2012). At first sight susceptibility to DCV in *Drosophila* would appear to be a clear example of this pattern, with a restriction factor called Pastrel explaining as much as 78% of the genetic variance in this trait (Cogni, et al. 2016). However, we have found that this belies considerable complexity within this locus. Strikingly, in a sample of just seven alleles from natural populations, we found four phenotypically distinct allelic classes conferring differing levels of resistance to DCV. Furthermore, both coding and *cis-*regulatory variants control resistance. The coding sequence variant that we characterised appears to be an old polymorphism that has been maintained at a relatively stable frequency, possibly as a result of balancing selection.

An amino acid polymorphism in Pastrel is the most important factor determining susceptibility to DCV. There are multiple lines of evidence to support this. First, this is the only genetic variant that can explain the largest changes in resistance that we see in two large genetic mapping experiments. Second, when populations have been artificially selected for DCV resistance, this site shows the largest increase in frequency in the entire genome (Martins, et al. 2014). Finally, when we modified this site in transgenic flies we verified that it is the cause of resistance.

The ancestral state at this site was the susceptible allele threonine. Three other major-effect polymorphisms that affect susceptibility to viruses in *Drosophila* have been identified at the molecular level, and in all cases the ancestral state was susceptible (Bangham, et al. 2008; Cao, et al. 2016; Magwire, et al. 2011). This fits with a model whereby genetic variation is arising because there is continual input of novel resistance alleles into populations from mutation, and these are then favoured by natural selection.

Resistance could evolve by improving existing antiviral defences or by altering the myriad of host factors hijacked by the virus for its own benefit. For example, in *Caenorhabditis elegans*, susceptibility to the Orsay virus is determined by a polymorphism that disables the antiviral RNAi defences (Ashe, et al. 2013), while bacteriophage resistance is frequently associated with changes to surface receptors used by the virus to enter cells (Longdon, et al. 2014). In a previous study we found that knocking down the susceptible allele of *pst* makes flies even more susceptible (Magwire, et al. 2012), while in this study we found that overexpressing the susceptible allele makes flies resistant. Therefore, the threonine to alanine mutation that we observe in *pst* is an improvement to an existing antiviral defence.

Patterns of genetic variation at the *pst* locus are complex. We found extensive structural variation, with multiple duplications and deletions of the gene present in natural populations. Furthermore, there is genetic variation in the expression of *pst*. There was a single QTL that controls *pst* expression, and this was centred on *pst* itself. Therefore, *cis* regulatory variants control *pst* expression.

Higher levels of *pst* expression are associated with increased resistance to DCV. This is unsurprising, as when we have experimentally altered *pst* expression by RNAi or by overexpressing the gene DCV resistance is altered. Both SNPs and structural variants in the region are associated with *pst* expression. However, the *cis*-regulatory variants which are causing increased expression could not be unambiguously identified because of linkage disequilibrium between these sites. Interestingly, the structural variants themselves were not significantly associated with survival after DCV infection, perhaps suggesting that they are not the main cause of variation in gene expression. Nonetheless, given the central role this gene plays in antiviral defence, it is tempting to speculate that these complex structural changes may have had some functional role, perhaps against other viruses (or we may simply lack the statistical power to detect effects on DCV) (Martins, et al. 2014).

Why is genetic variation in susceptibility to DCV maintained in populations? There is likely to be selection favouring alleles that increase resistance in natural populations because DCV is the most virulent virus that has been isolated from *Drosophila* and field studies have found it to be geographically widespread (Christian 1987) (although recent surveys have suggested that it may have a low prevalence (Webster, et al. 2015)). Pastrel has also been implicated in resistance to other viruses related to DCV (Martins, et al. 2014). Given that the resistant allele is likely to enjoy a selective advantage, an important question is why the susceptible alleles have not been eliminated by natural selection. To understand how selection has acted on the amino acid variant that causes resistance, we examined geographical variation in its frequency and patterns of linkage disequilibrium with neighbouring sites. We could detect no evidence of natural selection causing changes in allele frequency through time or space. This is in stark contrast to the partial selective sweeps that we have seen in the two other major-effect polymorphisms affecting virus resistance (Bangham, et al. 2008; Magwire, et al. 2011). These polymorphisms are in the genes *CHKov1* and *P62* (*ref(2)P*) and both confer resistance to the sigma virus. In both cases the resistant allele has recently arisen by mutation and has spread through *D. melanogaster* populations under strong directional selection. In comparison to these polymorphisms it is clear that the polymorphism in *pst* is relatively old and has been maintained at a stable frequency.

These population genetic patterns suggest that either the polymorphism has been evolving neutrally or it has been maintained by balancing selection, due to the benefits of resistance being balanced by harmful pleiotropic effects of the resistant allele on other traits. Long-term balancing selection can leave a signature of high divergence between the two alleles and elevated sequence polymorphism (Charlesworth 2006), but we have been unable to find any evidence of this in *pst*. This is not unexpected because the large effective population size of *D. melanogaster* means that linkage disequilibrium declines rapidly around *pst*, and this is expected to erode any signature of balancing selection (Charlesworth 2006). A very similar pattern of sequence variation was recently reported around a polymorphism in the antimicrobial peptide Diptericin that affects susceptibility to bacterial infection (Unckless, et al. 2016). This amino acid polymorphism is also found in the sibling species *D. simulans*, strongly suggesting it is maintained by balancing selection. Therefore, we cannot distinguish balancing selection and neutral evolution. While it seems likely that a polymorphism with such a large phenotypic effect is the target of natural selection, we would need data from natural populations to demonstrate that this was the case.

In *Drosophila* increased resistance against bacteria and parasitoid wasps is associated with reduced fecundity and larval survival (Kraaijeveld and Godfray 1997; McKean, et al. 2008). However, when populations of flies were selected for DCV resistance there was no detectable decline in other components of fitness (Faria, et al. 2015). Unfortunately, while it is clear the resistant allele of *pst* is not highly costly, this negative result is hard to interpret. First, if the benefits of DCV resistance in nature are small, then a small cost that cannot be detected in the lab will be sufficient to maintain the polymorphism. Without having an estimate of the harm flies suffer due to DCV infection in nature it becomes impossible to reject the hypothesis that the benefits of resistance are balanced by pleiotropic costs. Second, costs of resistance are typically only expressed in certain environments and may affect many different traits (Kraaijeveld and Godfray 1997; McKean, et al. 2008). It is possible that costs may not be detected if they are measured in the ‘wrong’ environment or the trait affected is not measured—for example, it may increase susceptibility to other pathogen genotypes.

The function and identity of viral restriction factors in invertebrates remains poorly understood, and the mechanism by which Pastrel protects flies against DCV is unknown. This contrasts with vertebrates where a diverse range of restriction factors has been characterised that inhibit all steps of viral infection (see (Yan and Chen 2012) for review). Studying natural variation in susceptibility to viral infection is proving a powerful way to identify novel restriction factors in *Drosophila* (Bangham, et al. 2008; Cao, et al. 2016; Magwire, et al. 2011), and future work on these proteins is likely to provide new insights into how invertebrates defend themselves against infection. One clue as to the function of Pastrel comes from its localisation to lipid droplets in the larval fat body (Beller, et al. 2006), as lipid droplets and lipid metabolism frequently play key roles in the viral replication cycle (Stapleford and Miller 2010). An alternative explanation is the reported involvement of Pastrel in the secretory pathway and Golgi organization (Bard, et al. 2006).

We conclude that a single gene, *pastrel*, is the dominant factor that determines the susceptibility of *D. melanogaster* to DCV. This is a complex locus, with multiple alleles conferring different levels of resistance, with polymorphisms affecting both the expression and protein sequence of Pastrel altering susceptibility to DCV. This gene has not been the target of strong directional selection, and the variation may be maintained by balancing selection. Overall, despite a single gene explaining most of the genetic variance in DCV susceptibility, this locus is remarkably complex.

## Acknowledgements

Thanks to Professor Jean-Luc Imler for helpful discussions, Yuk-Sang Chan for embryo microinjection. This work was supported by European Research Council (281668 to F.M.J) and Natural Environment Research Council ( NE/L004232/1 to FMJ).

